# Insights into the Structural Regulation of Polo-Like Kinase Activity using AlphaFold

**DOI:** 10.1101/2024.10.21.618045

**Authors:** Michael D. Wyatt, Campbell McInnes

**Author notes:** Corresponding Author Address: Drug Discovery and Biomedical Sciences, College of Pharmacy, 715 Sumter Street, University of South Carolina, Columbia SC 29208.

## Abstract

The Polo Like Kinases including the major family member, PLK1 are key regulatory enzymes controlling the cell cycle and mitosis. PLK1 is associated with poor survival rates in cancer and has been extensively investigated as an oncology drug target. Each member of the Polo like kinase family (PLKs 1-5) have two subdomains with independent functions and include the well conserved N-terminal kinase domain (KD) and the C-terminal polobox domain (PBD). The PBD is involved in the recognition of substrates primed by other kinases and in the PLK1 context is responsible for subcellular localization to specific sites in the nucleus including centrosomes and kinetochores. While the phosphosubstrate recognition site in PLKs 1-3 is highly conserved, its role in PLKs 2 and 3 is not well characterized and phosphopeptides that inhibit PLK1 have dramatically lower affinity for PLKs 2 and 3. An additional role of the PBD is its ability through domain-domain interactions with the KD to regulate PLK1 activity by an autoinhibited state of PLK1, conceptually similar to that which occurs through other kinases. Other mechanisms regulating PLK activity include the interchange between monomeric and dimeric forms, which inhibit or activate PLK1 during the cell cycle. Furthermore, PLK1 may exist as heterodimers with PLK2 and/or PLK3 and thus play context dependent roles. Here, through the use of the AlphaFold (AF) algorithm, structural insights into regulation of activity of the PLK1 and other family members have been obtained. These include dramatically different tertiary arrangement of the individual domains in each individual PLK. Analysis of the domain-domain interactions, interdomain and intradomain loops in each PLK sheds light onto plausible mechanisms by which the activity of each PLK is regulated and provides insights into the selectivity of phosphopeptides. The results also suggest a mechanism for the heterodimerization of PLK1 and PLK2 which has been observed in the literature.

## INTRODUCTION

Polo Like Kinases (PLKs) are Serine/Threonine kinases that regulate key processes involved in cell growth and division[1] with family members (PLK1-5) having individual expression levels, substrates, cell cycle functions, and sub-cellular location[2]. PLK1-3 have highly conserved kinase domains (KD, N-terminal) and polo box domain (PBD, C-terminal) but vary considerably in the length and composition of the interdomain connecting loop (Table 1)[2, 3]. As mutation of the IDL affects the activity of PLK1, its precise roles in the regulation of PLK1 and other PLK members remain to be determined[4]. PLK1 is the most extensively characterized in having its roles in mitosis (centrosome maturation, bipolar spindle formation, sister chromatin separation and cytokinesis among others) delineated[2] and has also been extensively pursued as a cancer drug target[3, 5]. Furthermore, numerous crystal structures have been solved for independent kinase and polo-box domains of PLK1 and have provided key insights into how substrates and inhibitors bind, thus supporting drug discovery efforts[6]. The structures of the KD of PLK1 have shed light into determinants for potent inhibition and show little variability in conformation when bound to various ATP competitive inhibitors[7, 8]. The PBD, which is essential for PLK1 localization and function, binds a phosphoserine/phosphothreonine (STP) motif, is critical for its sub-cellular localization and for substrate recognition prior to phosphorylating target substrates[9-11] has also been extensively characterized structurally as an independent domain. Greater variability of the PBD has been observed depending on the ligands and also in comparison to the autoinhibited structure of danio PLK1[12].

**Table 1.** Domain comparison of PLKs 1-5 (number in paraentheses are domain length)

|  | N-term | KD | IDL | PBD |
| --- | --- | --- | --- | --- |
| **PLK1** | 1-38 | 39-325 (287) | 326-364 (39) | 365-603 (238) |
| **PLK2** | 1-67 | 68-354 (287) | 355-445 (91) | 446-685 (239) |
| **PLK3** | 1-47 | 48-334 (287) | 335-423 (88) | 424-646 (223) |
| **PLK4** |  | 1-270 (270) | 271-594 (324) | 595-970 (376) |
| **PLK5** |  | 1-81 (81) | 82-150 (69) | 151-336 (186) |

While to date, over 3000 papers have been published on PLK1, far less is known about PLKs 2 (>300 publications) and 3(<300). As a result, they are less completely characterized in terms of their biological roles and structurally. From what is known, PLK2 has been shown to be a tumor suppressor kinase and has roles in synaptic plasticity, centriole duplication and regulating the transition between G1 and S phases of the cell cycle[13]. The role of PLK2 in synaptic plasticity and memory occurs through its regulation of the Ras and Rap signaling by phosphorylating RASGRF1 (Ras activator) and SIPA1L1(Rap inhibitor) leading to proteasomal degradation[14]. PLK2s role as a tumor suppressor is evidenced by its down-regulation in acute myeloid leukemias[15] and B-cell lymphomas[16]. Further evidence suggests that PLK2 down-regulation by miR-126 can lead to progression of leukemias[17]. PLK2 phosphorylation of CENPJ and NPM1is necessary for procentriole formation and centriole duplication[18] and its induction by p53/TP53 indicates participation in the mitotic checkpoint following stress[19].

PLK3 has roles characterized in cell cycle regulation, response to stress and Golgi disassembly. Known substrates of PLK3 include ATF2, BCL2L1, CDC25A, CDC25C, HIF1A, JUN, p53, PTEN and TOP2A[20-26]. PLK3 is required for entry into S phase and cytokinesis through its phosphorylation of BCL2L1 and has been shown to be involved in the DNA damage response by phosphorylating CDC25A, CHEK2, p53/TP53 and p73/TP73 in response to DNA damage, promoting the G2/M transition checkpoint and thus leading to inhibition of transcriptional activation and pro-apoptotic functions[27]. PLK3 is also involved in Golgi disassembly through a MEK1/MAP2K1-dependent pathway that induces Golgi fragmentation during mitosis[28].

Among this family, PLK4 diverges significantly in structure from the others. The sequence and furthermore the substrate specificity of PLK4 are variant from PLKs 1-3[29]. PLK4 localizes to the centrosome and controls appropriate centriole duplication prior to mitosis through recruitment of centrosome-related proteins including STIL[30] and SAS6[31-33]. Furthermore, PLK4 regulates centriolar satellite integrity and ciliogenesis, required for mitotic progression. Overexpression of PLK4 results in defects in centriole biogenesis, chromosome segregation, ultimately causing problems with cell cycle progression and potentially leading to tumorigenesis[34, 35]. Excessive PLK4 activity can lead to drug resistance to chemotherapy and contribute to metastasis, providing impetus for the development of inhibitors as anti-tumor therapeutics[29].

PLK5 remains the least-studied family member and less than 30 papers describing its study have been published. In contrast to the other family members, PLK5 does not seem to have a role in cell cycle progression probably because although its N-terminus has similarity to other PLKs, the kinase domain is nonfunctional due to truncation[36] and the PBD is not fully formed. PLK5 is mostly expressed in the brain where it modulates the formation of neuritic processes[37]. PLK5 gene silencing occurs brain tumors indicating that it plays a tumor suppressor role[37]. Further studies confirmed this in that upon examination of a wide variety tumors and comparison with normal tissues, results showed that PLK5 is significantly downregulated PLK5 expression in cancers and that expression remains consistently low in later stages indicating a greater role in tumor initiation[38].

AlphaFold is an artificial intelligence program that was recently developed by Google DeepMind. AlphaFold 2.0 was a dramatic breakthrough in its ability to essentially solve the protein folding problem computationally, can predict a protein’s 3D structure from its amino acid sequence with high accuracy and is very much competitive with experimentally determined structures. AlphaFold DB is a publicly accessible database of more than 200 million protein structure predictions from a vast array of organisms and species and has proved to be a tremendous resource to the scientific community. Here, AF structures of the PLK family are analyzed to provide comparisons into the conformational regulation of their activity (Figures 1-6). Furthermore, new AF predictions have been carried out on PLK mutants and to generate insights into the specific determinants required in for the regulatory architecture of each PLK.

**Figure 1.**
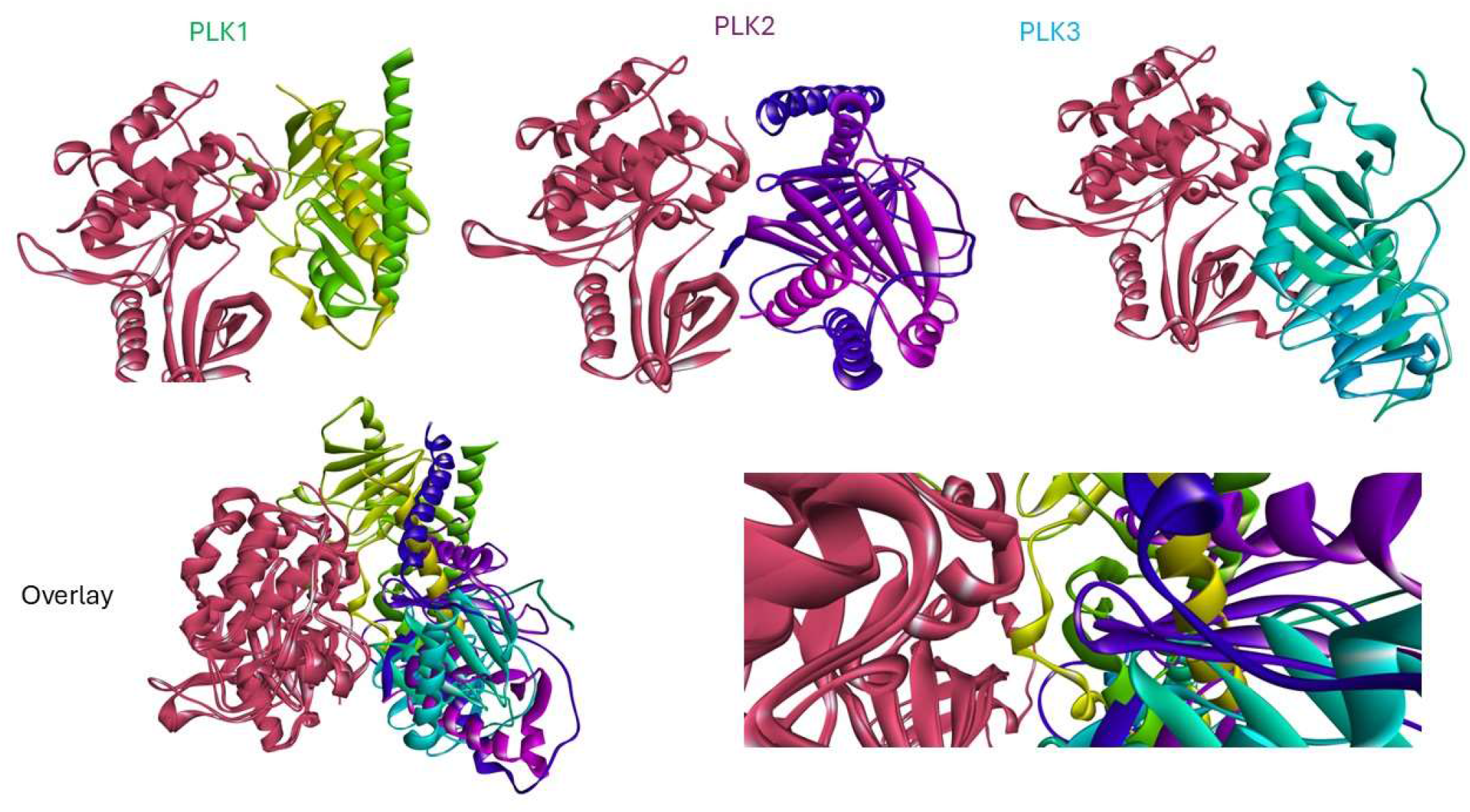
Comparison of AlphaFold models generated for PLKs 1-3. Individual PLK1s shown at the top of the figure are superimposed just using the KD residues to highlight the differences in the positions of the PBD in each case. The compact autoinhibited PLK1 structure is not observed in PLK2 or PLK3.

**Figure 2.**
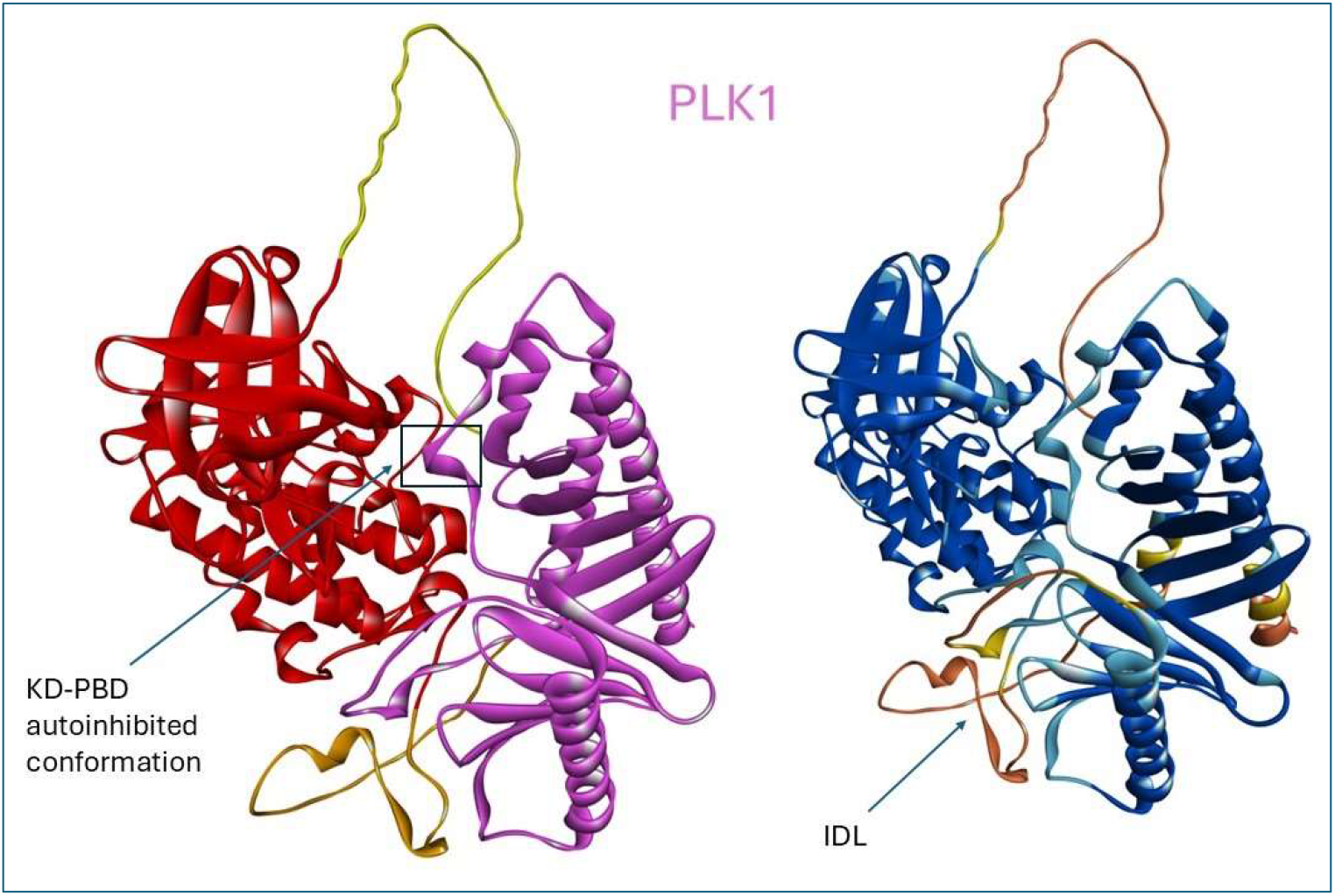
AlphaFold 2 Predicted structure of FL PLK1 (1-603). The left model is colored according to domain with the C-terminus in yellow (1-37), KD in red (38-325), the IDL in orange (326-364) and the PBD in magenta (365-603). The right model is colored according to pLDDT score: Dark blue - Very high (pLDDT > 90); light blue - High (90 > pLDDT > 70); Yellow-Low (70 > pLDDT > 50); Orange - Very low (pLDDT < 50).

**Figure 3.**
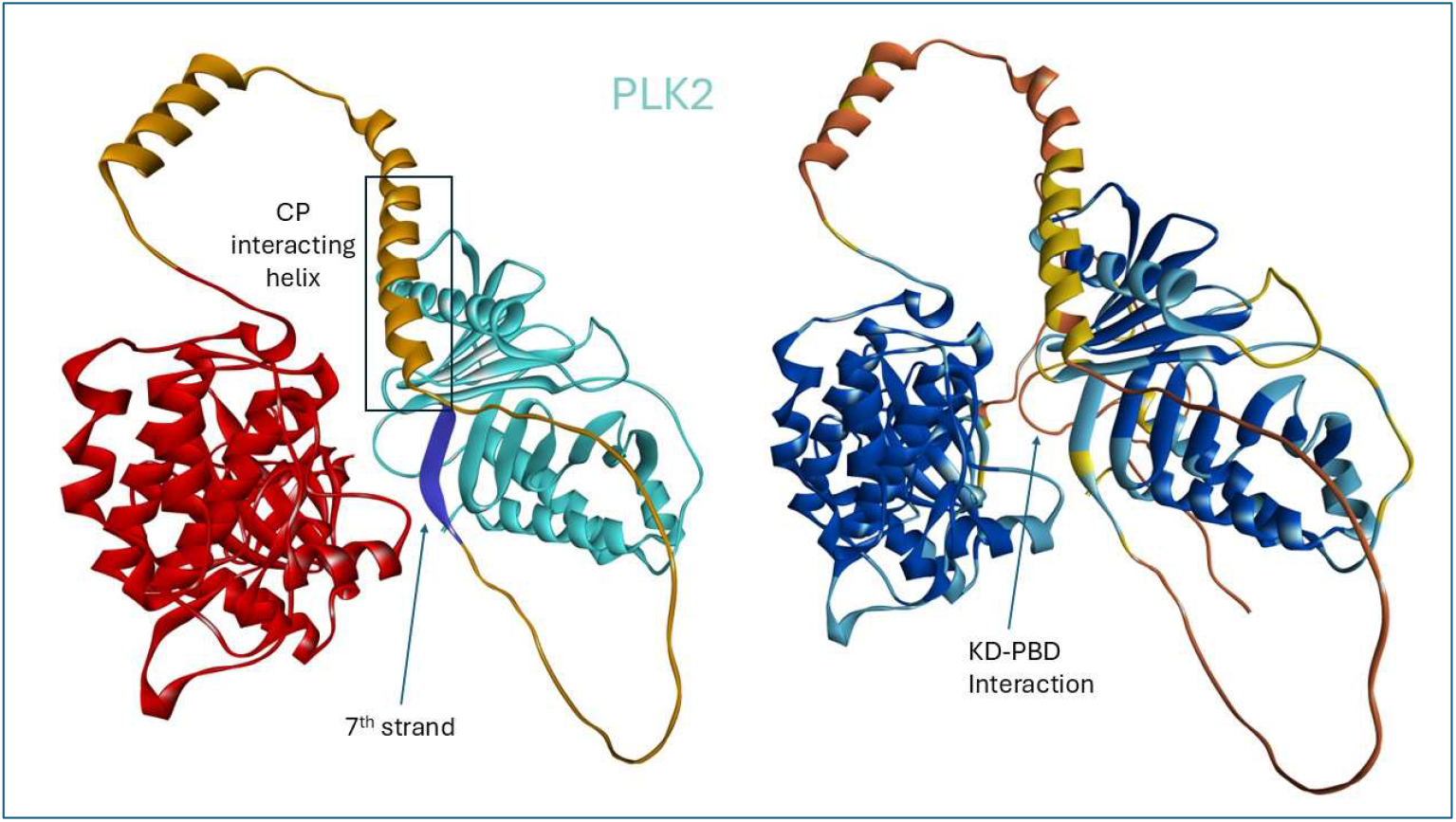
AlphaFold 2 Predicted structure of FL PLK2 (1-685). The left model is colored according to domain with the KD in red (68-354), the IDL in orange (355-445) and the PBD in magenta (446-685). The right model is colored according to pLDDT score: Dark blue - Very high (pLDDT > 90); light blue - High (90 > pLDDT > 70); Yellow-Low (70 > pLDDT > 50); Orange - Very low (pLDDT < 50).

**Figure 4.**
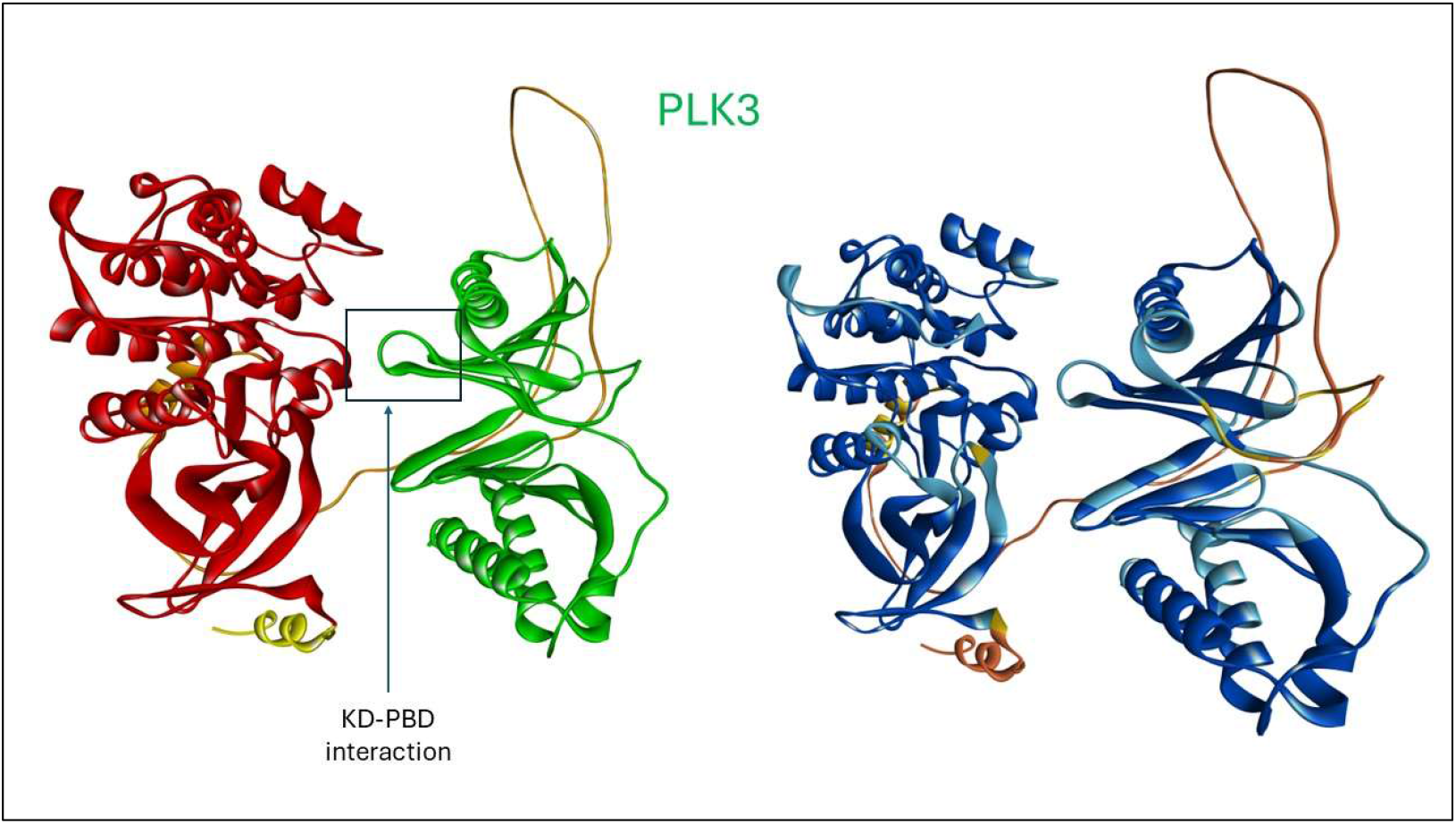
AlphaFold 2 Predicted structure of FL PLK3 (1-646). The left model is colored according to domain with the KD in red (48-334), the IDL in orange (335-423) and the PBD in magenta (424-646). The right model is colored according to pLDDT score: Dark blue - Very high (pLDDT > 90); light blue - High (90 > pLDDT > 70); Yellow-Low (70 > pLDDT > 50); Orange - Very low (pLDDT < 50).

**Figure 5.**
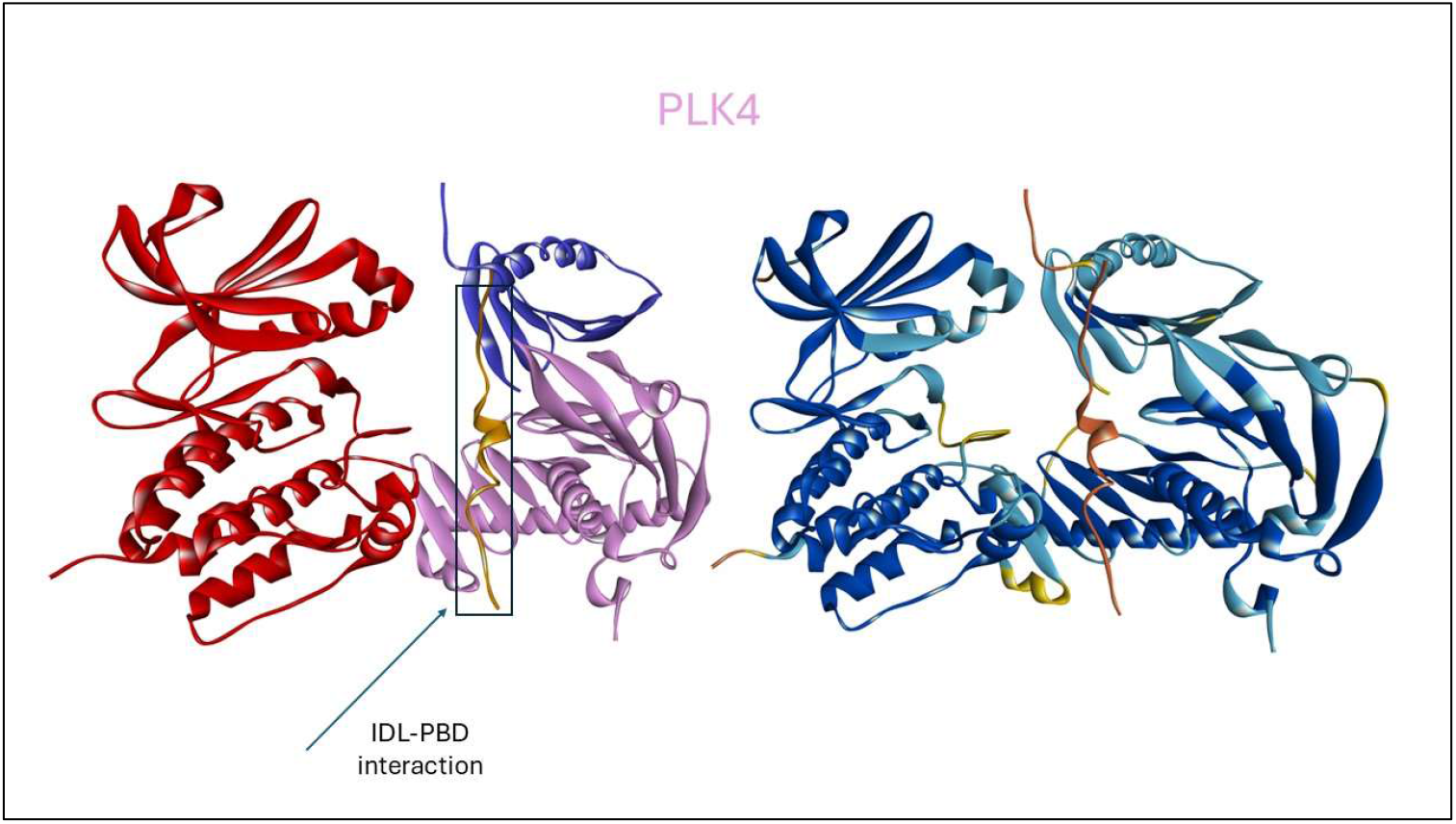
AlphaFold 2 Predicted structure of FL PLK4 (1-970). The left model is colored according to domain with the KD in red (1-270), the IDL in orange (467-486) and the PBD in magenta (595-970). The right model is colored according to pLDDT score: Dark blue - Very high (pLDDT > 90); light blue - High (90 > pLDDT > 70); Yellow-Low (70 > pLDDT > 50); Orange - Very low (pLDDT < 50).

## RESULTS

### Structural Insights into the Polo-Like Kinase family through AlphaFold

#### Polo-Like Kinase 1

While many structures of the independent kinase and polo-box domains exist, no experimental 3-D structures of FL PLK1 have been solved to date. While these structures have shed considerable light into the domain architecture and how ligands bind, it is likely that PLK1 exists in multiple conformational states and that flexible regions including the IDL and intradomain loops preclude crystallization. The first insights into the structural basis for how PLK1 is regulated were provided through a crystallization complex of the isolated domains (PDB ID: 4J7B)[4] using polo-like kinase KD and PBD from zebrafish (*Danio rerio*) stabilized by binding *of* a Map205 polypeptide (PBD-binding motif of microtubule-associated protein 205), a protein unique to *D. melanogaster*.

This ternary complex revealed that the KD is inhibited by the PBD with a large contact interface observed between the two domains. As this structure was somewhat artificial and in the presence of a non-mammalian inhibitory protein, a study of the AF generated full length human PLK1 3-D structure was undertaken. To provide further insights into human FL PLK1 activation, the structure of FL human PLK1 generated using AF[39-42] was analyzed. Analysis of the human FL PLK1 structure generated by AF in comparison with available crystal structures of individual KD and PBD domains revealed some remarkable observations. The AF structure shared high similarity with the Danio Rerio PLK1 structure (isolated KD and PBD co-crystallized [4]) but also provides insights that are absent in this artificial complex. As demonstrated by the Danio structure, the predicted structure was also found to be in the autoinhibited state (**Figure 6**) and shown to be essentially identical for the regions that are in common. These similarities confirm the regulatory mechanism in diverse homologues of PLK1, not surprising given the sequence identify between organisms is high. Furthermore, since PLK1 is known to adopt multiple conformational states and oligomeric forms dependent on the stage of the cell cycle and as AF generally generates low energy states of proteins, it is likely that the autoinhibited conformation is the most favorable state of PLK1 prior to post-translational modifications and binding of regulatory proteins such as Bora.

**Figure 6.**
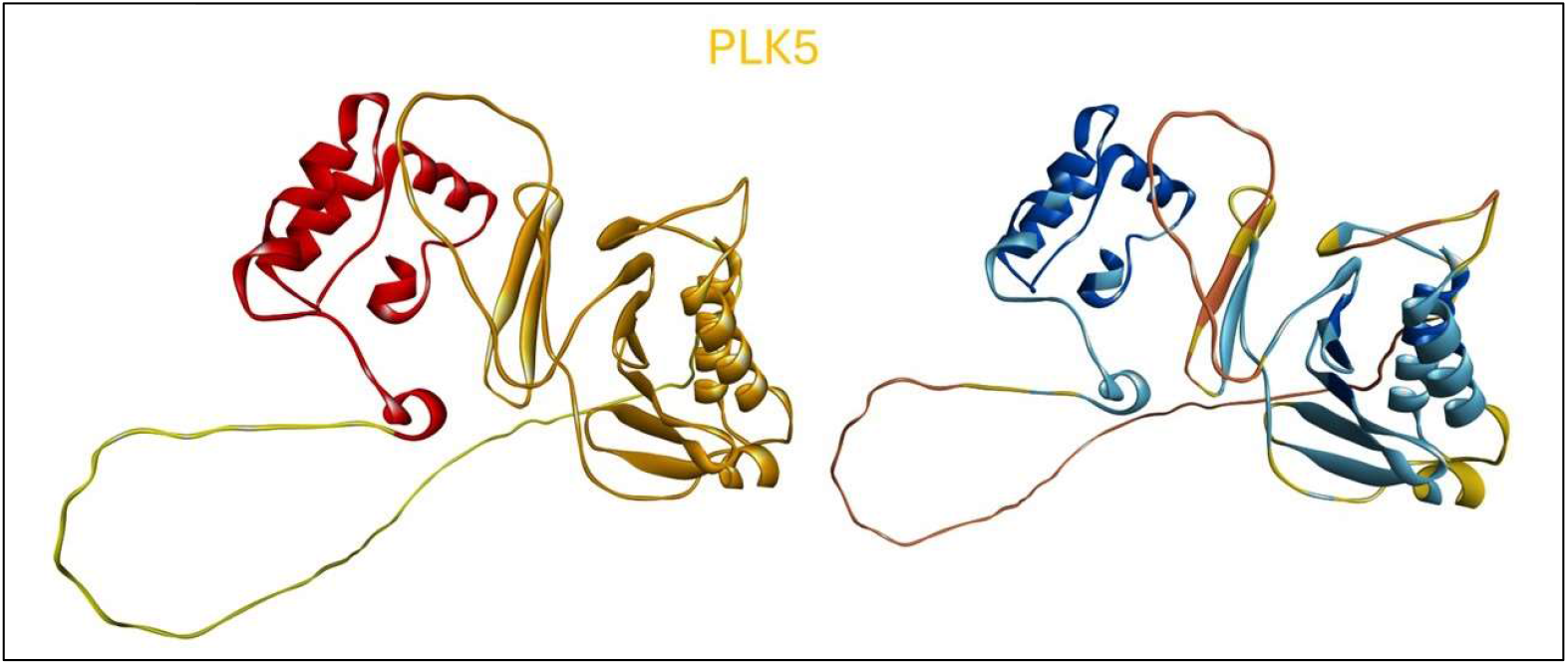
AlphaFold 2 Predicted structure of FL PLK5 (1-336). The left model is colored according to domain with the pseudo KD in red (1-81), the IDL in orange (82-150) and the PBD in magenta (150-336). The right model is colored according to pLDDT score: Dark blue - Very high (pLDDT > 90); light blue - High (90 > pLDDT > 70); Yellow- Low (70 > pLDDT > 50); Orange - Very low (pLDDT < 50).

The AlphaFold structure was further analyzed to provide insights into the missing regions of PLK1 that were either deleted in generating individual domains suitable for complex crystal formations or which were absent in electron density in the solved X-ray structure. One region absent in the Danio structure but which is known to play a key role in regulation of PLK1 activity is the roughly 50 residue sequence (residues 315-365) spanning the sequence connecting the end of the structured kinase domain with the beginning of the 3-D fold of the PBD known as the polo-cap. This sequence comprises the Interdomain Connecting Loop (IDL) and in addition to its role acting as a hinge allowing conformational opening of the two domains. It also contains the destruction box (DBOX, R337) which facilitates the degradation of PLK1 through ubiquitination. The AlphaFold structure suggests that the IDL is partially structured, contains a one helical turn and also confirms that the α-helix of the PC can be extended almost two more turns and covers the sequence (365-387, equivalent residues 365-371 missing in 47B). Furthermore, it provides insights into the missing segment in the danio complex from residues 489 to 498 which comprises most of the residues of the linker 2 region (L2 loop discussed further below). L490 of PLK1 makes a number of contacts to M486 and Y485, residues that are part of the cryptic or Tyr pocket, an important determinant of PBD substrate recognition.

Comparison of the peptide-bound, isolated PBD domain structures with the FL PLK1 AF structure yielded information not observed in the Danio autoinhibited PLK1 [12]. Although the PBD structures are similar, the linker 2 region (L2 – human residues 490-502), differs between the two structures. L2 C-terminal to PB1 diverges significantly in the AF structure and it is this region that contacts residues of PBD substrates that are C-terminal to the pThr. The isolated PBD bound to the peptide possibly represents the conformation of L2 that exists in the open conformation of FL PLK1.

A detailed analysis of the stabilizing interactions between the two domains in the autoinhibited conformation of PLK1 was carried out (Table 1). An extensive network of hydrophobic interdomain contacts is present suggesting that the autoinhibited state is highly stable. The PBD residues contributing to the closed conformation include the 403-407 loop of PB1 that is in close proximity to the ATP binding site of the KD and contacts L59, R136, E140 and R144. I408 and F409 make extensive contacts to the C-terminal region of the KD including residues I316, T317, L319 and T320. Residues of the L2 loop (500-502, especially R500) and L505 of PB2 make extensive contacts to these same residues of the KD but also to P323, R324 and F325. Of all the PBD residues, L505 makes the most extensive contacts to the KD and therefore contributes the most to stabilizing the closed conformation. Residues 506-509 make several interdomain contacts to the KD although individually not contributing as much as L505. Additional residues from PB2 that contribute to stabilization of the autoinhibited state include L546 and M547 which contact residues 140-146 of the KD and are close in proximity to the P403 loop that is closest to the ATP binding site.

#### Polo-Like Kinase 2

The full-length structure of PLK2 predicted by AF was analyzed and furthermore compared to the structural information currently available on the isolated kinase[43] and polo-box domains[44]. Three experimental structures have been solved to date and these include one for the KD[43] and two for the PBD[44, 45]. Superposition of the KD with AF structure revealed that the two are essentially identical. Comparison of the PBD structures with the AF predicted conformation revealed that the domains are very similar also. Comparison of the PLK1 and PLK2 AF structures however revealed dramatic differences with the most significant one being that PLK2 does not adopt a highly compact autoinhibited structure and in contrast to PLK1 there are few intra domain (PBD-KD) contacts (table 2). Comparison of the corresponding PLK2 residues that make stabilizing interactions in PLK1 reveal that these are not conserved between the two family members. While PLK1 has a large network of hydrophobic interdomain contacts leading to the closed conformation, PLK2 has few interactions between the domains and are mostly between polar side chains. Comparison of the residues that stabilize the PLK1 closed conformation with those found in PLK2 was carried out. While I408 of PLK1 is semi conserved as a F501 in PLK2, F409 is changed to Q502. L505, L546 and M547 are critical for stabilizing PLK1 in the autoinhibited state and their counterparts in PLK2 are R597, E639 and E640 respectively. Analysis of these differences alone strongly suggests that a similar autoinhibited state for PLK2 would not be favorable.

**Table 2.**
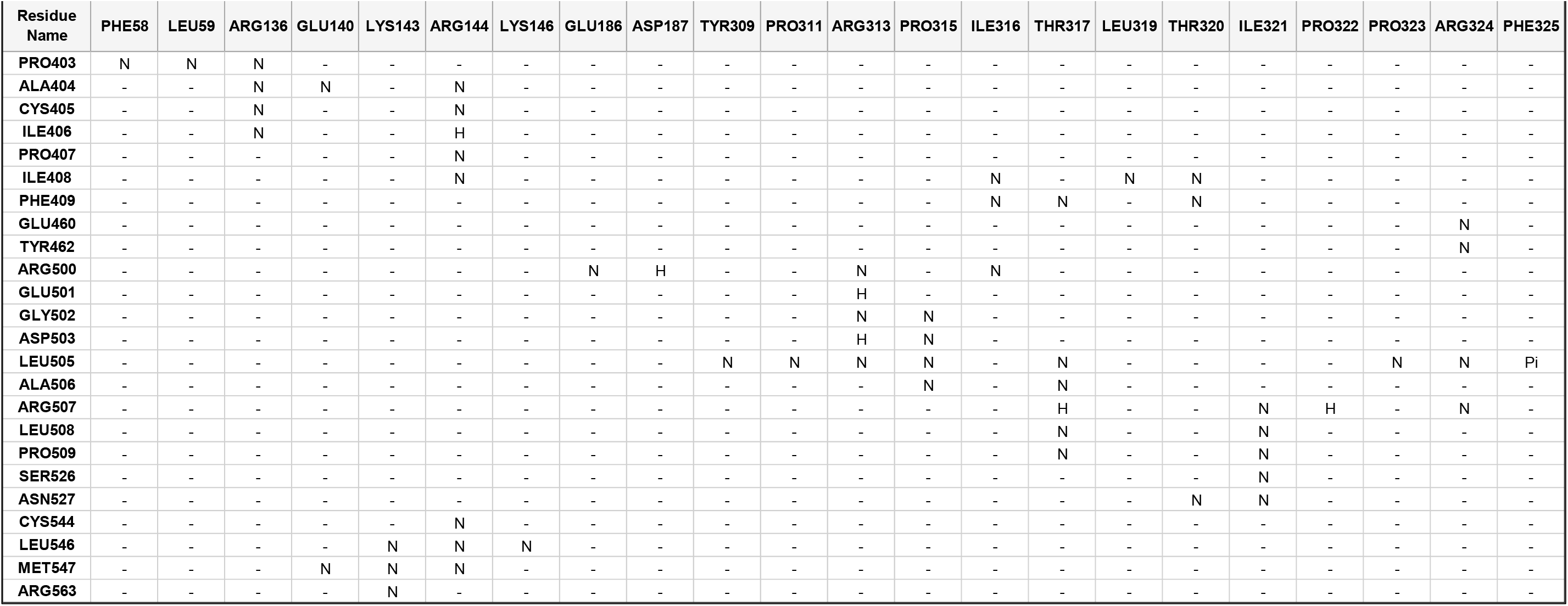
Interactions between the KD and PBD in PLK1 Analysis of the KD-PBD domain-domain interactions in the autoinhibited conformation of PLK1 generated with AF. An extensive network of stabilizing interactions is observed between the two domains (N= nonpolar, H = H bond), Pi interactions), SB = salt bridge)

**Table 3A, B.**
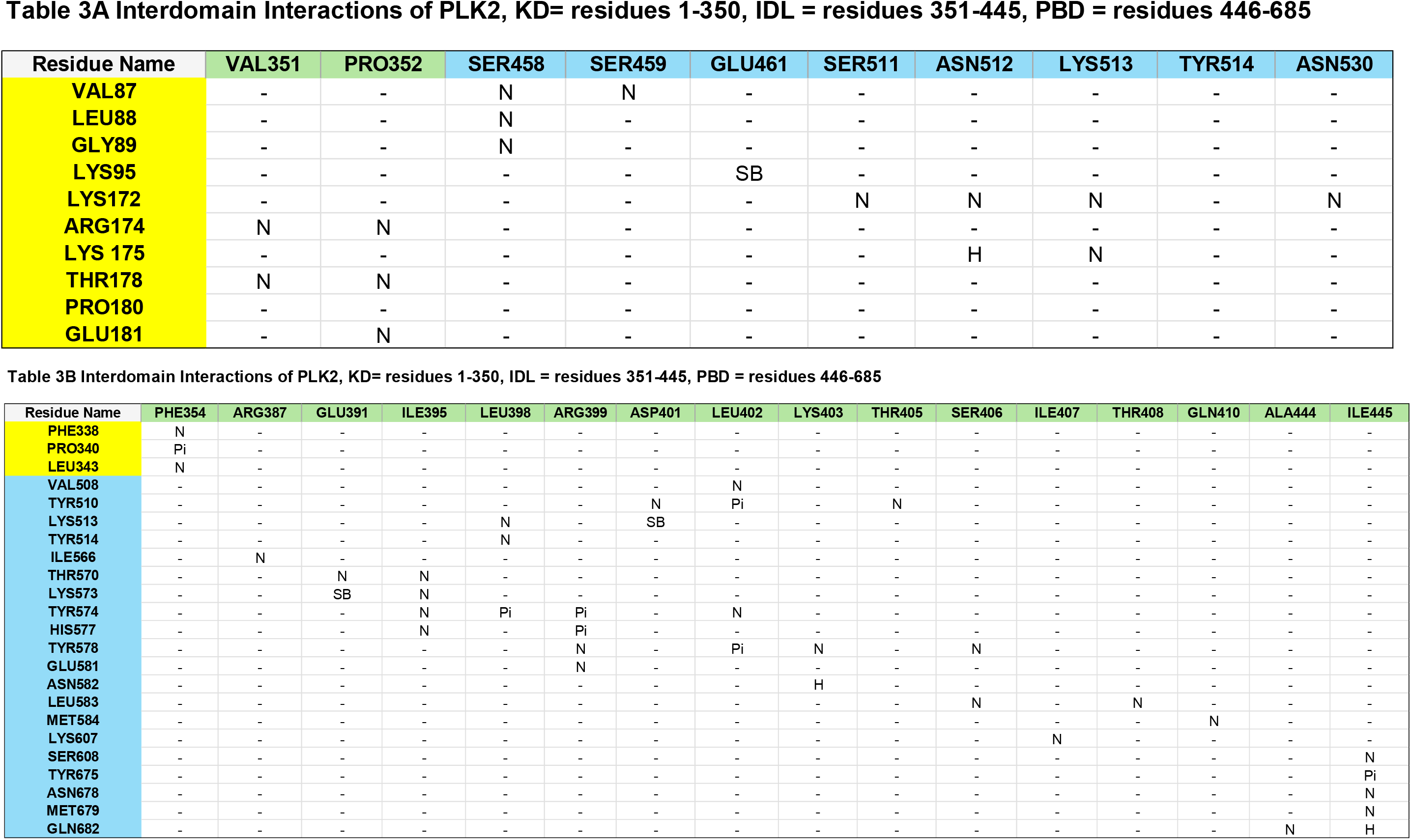
Analysis of the KD-PBD domain-domain interactions in the autoinhibited conformation of PLK2 generated with AlphaFold (N= nonpolar, H = H-bond, SB = salt bridge). KD, IDL and PBD residues are highlighted in yellow, green and blue respectively.

**Table 4.** Interdomain Interactions of PLK3 Analysis of the KD-PBD domain-domain interactions in the autoinhibited conformation of PLK3 generated with AF (N= nonpolar, H = H-bond, SB = salt bridge).

| Residue Name | LYS70 | ARG75 | GLU261 | THR262 | ALA263 |
| --- | --- | --- | --- | --- | --- |
| LYS473 | - | - | H | H | N |
| LYS568 | N | - | - | - | - |
| THR569 | N | N | - | - | - |
| ASP570 | N | N | - | - | - |

**Table 5.**
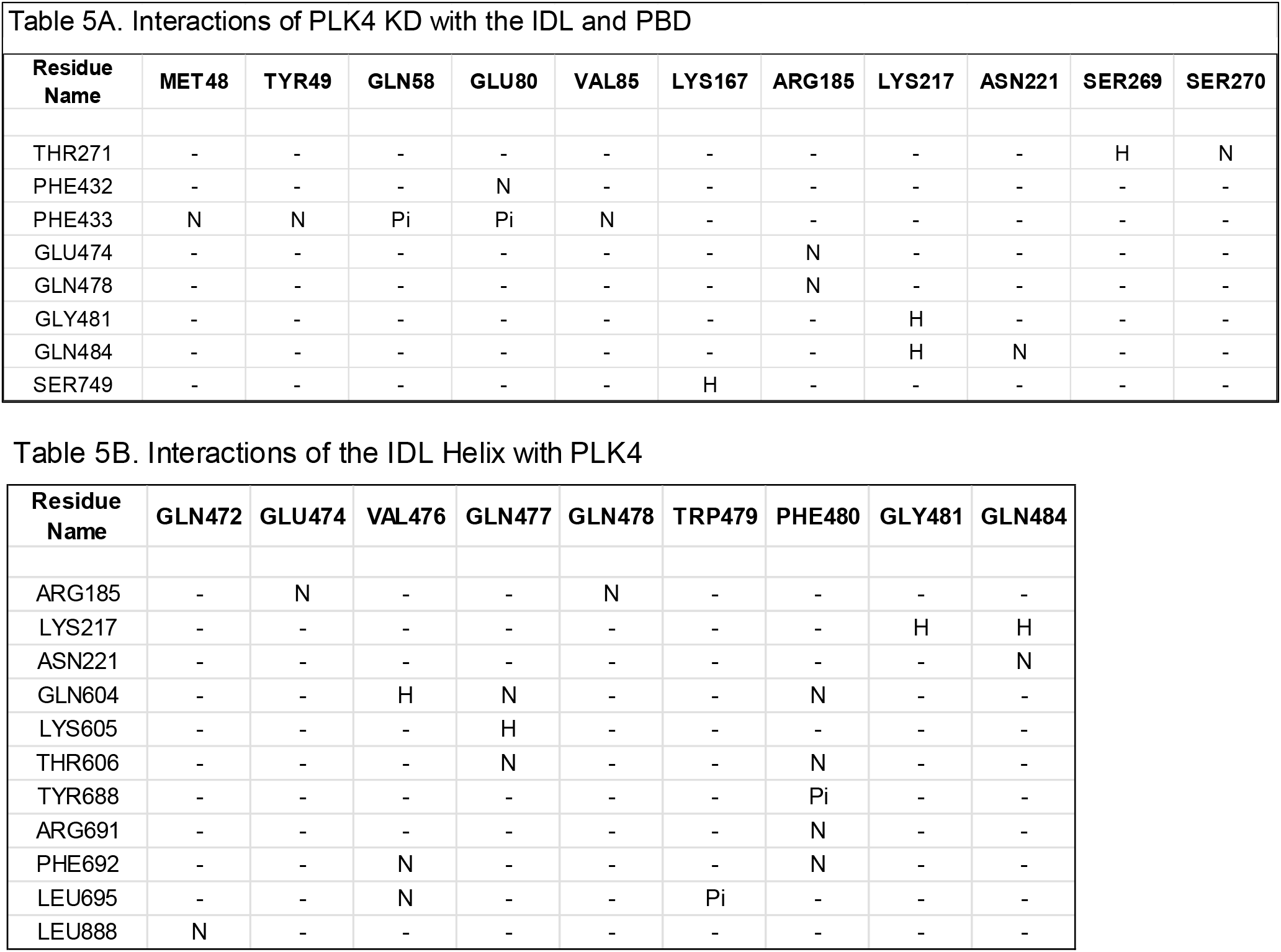
Analysis of the KD-PBD domain-domain interactions in the autoinhibited conformation of PLK4 generated with AF (N= nonpolar, H = H-bond, SB = salt.

Furthermore, the PLK2 AF structure revealed several secondary structure elements that are not evident from the isolated KD or PBD crystal structures. Relative to PLK1, it has a much larger IDL consisting of residues 350-454 and is predicted to contain two helical segments and a β-strand which forms the 7^th^ strand of a 6 strand anti-parallel β-sheet. Furthermore, the loop connecting this strand with the first helix of the PBD possesses a salt bridge between E461 and K95 of the KD, bringing this loop in close proximity to the ATP binding site of the KD of PLK2 It does not however block access to the extent that the P403 containing loop interacts with the KD in PLK1.This may also be a result of the dramatically longer inter domain connecting loop in PLK2. In fact, this larger loop seems to have a propensity to form secondary structure based upon the AF prediction. There are two helical segments in the PLK2 IDL from residues 358 to 373 and from 377 to 404. An interesting observation is that the longer of these two helical segments (377-404) makes significant interactions with the PBD and binds into the cryptic or tyrosine pocket that has shown to be a key interactor with certain substrates of PLK1 and furthermore which can be engaged in the design of potent inhibitors of the PBD. Key interacting residues of the IDL with the PBD include E391, L398, R399, L402 and K403. L398 and L402 engage Y574, F575, and Y578 of the PLK1 PBD cryptic pocket. Further insights into the secondary and tertiary structure of PLK2 were provided from an additional region of the IDL that is absent in the isolated domain of the PLK2 PBD. While residues 405-446 form a large unstructured loop, 447-453 form an additional and 7^th^ β-strand with the 6-strand antiparallel contiguous sequence of the PBD β6α motif between 602 and 656. Residues 454-465 comprise a loop structure which then connects with the 466-486 α-helix. E461 is engaged in a salt bridge with K95 bringing this loop in close proximity to the ATP binding site of the KD of PLK2 although does not block access to the same degree that the P403 containing loop interacts with the KD in PLK1. When the BI2536/PLK2 KD structure is superimposed with the AF structure there is overlap of the inhibitor with a section of this loop containing S457 suggesting it would act to displace the interdomain interactions in an analogous fashion to what is observed in the PLK1 context. Comparisons with of the PLK2 AF structure with two independent crystal structures of the isolated PBDs which exist in dimeric form in the unit cell. Superposition of the two monomers reveals that two very different conformations of the L2 loop exist in each one, where one is much more open and exposed than the other one. The AF structure indicates that the more compact L2 loop conformation exists in the full-length context. This is consistent with the observation that the open L2 structure exists as it interacts with another PBD in the asymmetric unit. Interestingly this interaction consists of phosphosubstrate like interactions from D588 of forming electrostatic interactions with K631and R650 and M584 perfectly docks into the cryptic hydrophobic pocket formed by Y510, Y574, and Y578, two of which are mentioned above as interacting with the longer helix of the IDL. This phosphosubstrate mimicking sequence intriguingly suggests that this might be a relevant mechanism of PLK2 dimerization with another PLK2 molecule or for heterodimerization with PLK1, for which evidence exists[46]. In addition, the cryptic pocket engaging helix present in the IDL of PLK2 could allow heterodimerization with PLK1 since this pocket is highly conserved between PLKs 1-3.

#### Polo-Like Kinase 3

To date, the only one crystal structure containing a PLK3 sequence available is that of the KD from residues 52-332 and bound to 4-[[(4R)-5-cyclopentyl-4-ethyl-3a,4-dihydro-3H-[1,2,4] triazolo [4,3-f]pteridin-7-yl] amino] -N-cyclopropyl -3-methoxy-benzamide in the catalytic cleft (4B6L, unpublished). To gain further insights into FL PLK3, the AF predicted structure was analyzed. Close examination of the AF structure for PLK3 indicates that it also does not form the compact autoinhibited structure found in PLK1. Furthermore, compared with PLK2, there are even fewer stabilizing interactions between the two domains. Only a few H-bond and non-polar contacts are observed between the KD and PBD with K70 and R75 of the KD interacting with K568, T569 and D570 and E261 of the PBD. Further interactions include T262 and A263 of the KD contacting K473 of the PBD. The PBD loop containing K473, in a similar fashion to PLK2, interacts in the proximity of the ATP binding site but only partially occludes it. Analysis of the PLK3 counterparts of essential residues stabilizing the closed conformation of PLK1 indicates that like with PLK2, these are not conserved with the counterpart of I508 being L461 in PLK3 while F509 is V462 in PLK3. L505, L546 and M547 of PLK1 are critical for stabilizing the autoinhibited state and their counterparts in PLK3 are V557, E599 and P600 respectively. The IDL of PLK3 is of similar length to that of PLK2 and covers residues 330-426 however has much less secondary structure according to the AF prediction. Only one helical segment is present covering residues 338-353 and the second helix contacting the PBD cryptic pocket and the 6^th^ β-strand are not predicted to occur.

#### Polo-Like Kinase 4

As mentioned, PLK4 diverges significantly from PLKs 1-3 not only in terms of the sequence homology of the kinase and polo-box domains but also in that the IDL is dramatically longer than even that of PLKs 2 and 3[2]. Numerous crystal structures have been solved for the KD[47] and for the cryptic PB[48] and PB3[49] regions of the C-terminal domain but again no full length structure exists. The KD is from residues 1-270 and has 42% identity with the PLK2 KD and 40% with the PLK1 KD. In contrast, the PLK2 KD has 52% identity with that of PLK1. The IDL from 271-594 is thus over 300 residues compared to 99 with PLK2 and 88 with PLK3. While most of the IDL is predicted to be intrinsically disordered, residues 467-486 form an extended structure with an α-helical segment in the center. While the overall prediction confidence level of the IDL is low, an interesting observation nonetheless is that the extended and helical segments are predicted to insert between the KD and the PBD and form plausible stabilizing interactions with each domain. Of particular note, are the multiple stabilizing contacts between F433 and M48, Y49, Q58, E80 and V85. Other important interactions are those of V476, Q477 and F480 of the helical segment and 604-606, 688-695. The PBD of PLK4 is from residues 595-970 and contains 3 polo boxes. The two PBs in the N-terminal region of the PBD are from 595-815 and this region overall is known as the cryptic polobox. Despite structural similarity, the sequence similarity with the PBDs of PLK1-3 is minimal and less than 30%. Of this cryptic PB, residues 700-815 are structurally very similar to the PBD of PLKs 1-3 but lacking the 2^nd^ β6α motif formed by residues 408-509 of PLK1. The remaining C-terminal sequence of PLK4 contains a disordered loop region between residues 816-885 and the structured domain from 886-970 known as PB3. This domain is required for interactions with PLK4 substrates and regulatory proteins such as CEP152, which form a complex and is required for centrosomal duplication. Again, while structural similarity exists PB3 has a unique sequence with no appreciable identity to any other protein (<30%).

#### Polo-Like Kinase 5

Of all the PLKs, PLK5 diverges the most in terms of sequence, structure and function. It has 336 residues, only half the number of PLK2. As mentioned, PLK5 does not have a functional KD as it is severely truncated compared to the other PLKs and although the identify is about 52% for PLK2 and PLK3 and 45% for PLK1, the homologous region only consists of about 80 residues (1-81) corresponding to the C-lobe of the KD of other PLKs. The IDL of PLK5 from residues 82 to 150, is unstructured in the AlphaFold model and the remainder of the sequence corresponds to a truncated PBD. Its homology is low compared to the other PBDs with 25, 27 and 32% identity for PLK1, 2 and 3 respectively. An interesting observation is that the C-terminus of the IDL (residues 152-159) forms a 7^th^ antiparallel β-strand with the β6α motif of PB2 somewhat analogous to that observed with PLK2. The polocap helix is intact, present from residues 170-185 and is similar to that of the other PLKs especially 2 and 3. Interestingly, it is truncated in the internal sequence of the PBD. The most identical region is residues 200-225, an antiparallel β-strand, and which contains many of the important phosphosubstrate recognition site in PLK1 and includes essential residues including W414 and Y417, Y421 and Y425 (comprise a significant part of the cryptic pocket). Following this, a large deletion in PLK5 results in loss of the helical segment and three of the six β-strands of the β6α motif of PB1 found in PLKs 1-3. This region contains the residues of PLK1 essential for stabilizing the autoinhibited structure of PLK1 identified in the Danio and human AF PLK1 structure. Following this, the fold of PB2 is almost identical to PLK1 although the key HTK phosphate binding residues are not conserved in PLK5 confirming its inability to recognize substrates of PLK1 and indicating that it does not participate in sub cellular localization in the same way.

## CONCLUSIONS

A detailed structural analysis of AlphaFold generated structures of the Polo-like kinase family has been carried out. This analysis provides novel insights into the conformations that each PLK member adopts and therefore potentially how they are regulated. The major conclusions of this study are that human PLK1 adopts a very similar autoinhibited conformation already identified in the artificial complex solved using independent PLK1 domains from Danio and stabilized by binding of the drosophila MAP205 peptide. In addition, the PLK1 AF structure provides additional information on the missing portions of PLK1 including the interdomain connecting loop and the C-terminal region of the Polo-Cap. Regarding PLK2 and PLK3, AF structural analysis and comparison of the residues responsible for stabilizing the closed conformation observed in PLK1, strongly suggests that these two family members do not adopt the stable autoinhibited structure found in PLK1. The network of hydrophobic contacts observed to stabilize the closed PLK1 structure is not present as these residues are primarily polar. PLK2 and 3 have relatively few interdomain (KD-PBD) contacts in contrast to PLK1. An interesting observation with PLK2 is that the considerably longer IDL (compared to PLK1), has a significant propensity to form α-helical structure. One of these helices makes numerous contacts with the cryptic pocket of PLK2, adjacent to the phosphopeptide recognition site. It inserts hydrophobic side chains into this pocket in a similar fashion to that observed in the MAP205/PBD interactions observed in the artificial danio complex with PLK1. As the cryptic pocket is essential for substrate interaction, in the PLK1 context, it is possible that this represents a type of autoinhibition of PLK2. Furthermore, as the cryptic pockets are conserved in PLKs1-3, a plausible scenario is that PLK2 could form heterodimers with PLK1 and PLK3, as there is evidence that this plays a role in desensitizing PLK1 to ATP competitive inhibition[46]. The AF models for PLK4 and 5 are less insightful due to the sequence and structural divergence of these members relative to PLK1-3 and the low confidence levels of the full-length structure. While little functional relevance can be concluded from these, insights into the interdomain interactions for PLK4 and how the truncated domains affect the folding of the KD and PBD in PLK5 have been obtained and their implications discussed. Analysis of these models and the conclusions drawn, postulate several lines of investigation to probe the conformational regulation especially of PLKs 1-3. Such studies will enable delineation of roles of the individual domains and connecting loops in the functional activity including that of homo and heterodimerization shown to play key roles in PLK activation.

## EXPERIMENTAL

PLK structures were downloaded from the AlphaFold database and were analyzed using Biovia Discovery Studio 2024 Visualizer. Interdomain interactions show in Tables 2-6 were generated using the interaction analysis module of DS2024. Some structures were recalculated using AlphaFold 3 which is freely accessible through the DeepMind website. Conclusions were not significantly affected after analysis of the AF3 structures.

